# Homeostatic mechanisms may shape the type and duration of oscillatory modulation

**DOI:** 10.1101/615450

**Authors:** Erik J. Peterson, Bradley Voytek

## Abstract

Neural oscillations are observed ubiquitously in the mammalian brain, but their stability is known to be rather variable. Some oscillations are *tonic* and last for seconds or even minutes. Other oscillations appear as unstable *bursts*. Likewise, some oscillations rely on excitatory AMPAergic synapses, but others are GABAergic and inhibitory. Why this diversity exists is not clear. We hypothesized Ca ^2+^ -dependent homeostasis could be important in finding an explanation. We tested this hypothesis in a highly simplified model of hippocampal neurons. In this model homeostasis profoundly alters the modulatory effect of neural oscillations. Under homeostasis, tonic AMPAergic oscillations actually *decrease* excitability and *desynchronize* firing. Tonic oscillations that are synaptically GABAergic–like those in real hippocampus–don’t provoke a homeostatic response, however. If our simple model is correct, homeostasis can explain why the theta rhythm in the hippocampus is synaptically inhibitory: GABA has little to no intrinsic homeostatic response, and so can preserve the pyramidal cell’s natural dynamic range. Based on these results we can also speculate that homeostasis may explain why AMPAergic oscillations in cortex, and hippocampus, often appear as bursts. Bursts do not interact with the slow homeostatic time constant, and so retain their normal excitatory effect.

**New and Noteworthy:** The intricate interplay of neuromodulators, like acetylcholine, with homeostasis is well known. The interplay between oscillatory modulation and homeostasis is not. We studied oscillatory modulation and homeostasis for the first time using a simplified model of hippocampus. We report a paradoxical result: Ca-mediated homeostasis causes AMPAergic oscillations to become effectively inhibitory. This result, along with other new observations, means homeostasis might be just as complex and important for oscillations as it is for other neuromodulators.

## Introduction

Neuromodulation and homeostasis can be thought of as inter-linked but opposing phenomena. Neuromodulation perturbs excitability, which we define as the propensity for a stimulus to elicit an action potential. Homeostasis acts to quench this perturbation, driving the excitability of the cell back to a biologically desirable set point (LeMasson, Marder, and Abbott 1993; Abbott and LeMasson 1993). While the interplay between chemical modulators and homeostasis has been studied for more than 20 years (LeMasson, Marder, and Abbott 1993; Abbott and LeMasson 1993; Golowasch et al. 1999; Marder, O’Leary, and Shruti 2014; Marder, Goeritz, and Otopalik 2015; Gutierrez, O’Leary, and Marder 2013), the relationship between population-level synaptic modulations–like neural oscillations–and homeostasis is not well understood. It is not well understood theoretically, or well-studied experimentally.

Like neuromodulators, neural oscillations are ubiquitous and alter the excitability and firing statistics of cells throughout the brain. Uniquely though, oscillations group action potentials into synchronous windows of activity (Lisman and Jensen 2013; Voytek and Knight 2015). Grouping action potentials improves their signal to noise, increases the number of coincident firing events (Chen et al. 2013; Zhou et al. 2015; Voytek et al. 2015; Peterson and Voytek 2017), that in turn drives learning at individual synapses (Muller, Brette, and Gutkin 2011; Song, Miller, and Abbott 2000; Markram 1997).

We propose the same mechanisms that link homeostasis with chemical neuromodulation should also come into play with oscillatory modulation. After all, both kinds of modulation can lead to tonic changes in spiking and, as a result, changes in Ca^2+^ concentration (Liu et al. 1998). To test this, we model activity-dependent intrinsic homeostasis in a feed-forward population of hippocampal pyramidal cells (Siegel, Marder, and Abbott 1994). By feed-forward we mean there are no lateral or recurrent connections in the population. Firing activity is driven only by afferent synaptic input, and homeostasis is mediated by a Ca^2+^-dependent mechanism (Golowasch et al. 1999; Marder, O’Leary, and Shruti 2014; Marder, Goeritz, and Otopalik 2015; Gutierrez, O’Leary, and Marder 2013; O’Leary et al. 2014) whose early work was pioneered by LeMasson (LeMasson, Marder, and Abbott 1993; Abbott and LeMasson 1993). In our model Ca^2+^ acts as a sensor for tonic changes in the membrane voltage. To counter tonic changes in Ca^2+^ levels, the expression of ion channels is altered, returning the Ca^2+^ level to a predefined “good” value (Golowasch et al. 1999; O’Leary et al. 2013). Following Siegel (Siegel, Marder, and Abbott 1994), increases in Ca^2+^ lead to downregulation of Na^+^ and upregulation of fast K^+^ channels. To minimize the effect of homeostasis on overall action potential magnitude, we also added a KCa channel (Siegel, Marder, and Abbott 1994). In real cells intrinsic homeostasis changes the expression level of ion channels (O’Leary et al. 2013). Here, these details are not directly simulated. Instead we mimic the net or bulk effect of all ion channels using a single equation (LeMasson, Marder, and Abbott 1993; O’Leary et al. 2013; O’Leary et al. 2014).

Given that chemical modulators operate on both short and long times scales (Marinelli and McCutcheon 2014; Marder, O’Leary, and Shruti 2014; Cohen, Amoroso, and Uchida 2015; Daw, Kakade, and Dayan 2002), we examined two timescales of oscillatory modulation. We contrast the effects of short 3-cycle bursts of oscillation to long lasting tonic rhythms. Both are observed *in vivo*, and are known to play different physiological, cognitive, and computational roles (Lundqvist et al. 2016; van Ede et al. 2018a; Peterson and Voytek 2017). We also explore synapse type, examining both AMPA- or GABA-ergic oscillations.

## Results

We assessed the impact of homeostasis in a simple model that received oscillatory inputs that are either transiently bursting, or long lasting and tonic. We found that increasing the strength of AMPAergic oscillatory inputs resulted in a homeostatic response that, when reaching an appropriate level of average calcium, overcompensates to actually decrease the postsynaptic neuron’s spiking. That is, paradoxically, an increase in drive– when combined with homeostasis–results in a decreased response.

Specifically, we study Ca^2+^-mediated homeostasis in a feed-forward population of 100 Hodgkin-Huxley neurons. By feedforward we mean neurons in the population are driven only by the inputs to the model. Cells have no lateral or recurrent connections. We modulate this population with neural oscillations. Oscillatory input for each cell in Hodgkin-Huxley population was modeled as a set of about *M* = 100 Poisson neurons (Figure 1**b**-**c**). With a probability of 0.1 these were randomly drawn from a pool of 1000 Poisson neurons, each of which was independently simulated. To change the level of oscillatory modulation strength, we altered the firing rate of the entire Poisson population. Average synaptic strengths were fixed, but drawn from a uniform distribution for each simulation (see *Methods*). An oscillation’s temporal duration was simulated as either tonic (Figure 1**b**, top), which lasted for the entire experiment (20 sec), or as a 3-cycle burst (Figure 1**b**, bottom). Bursts were delivered only at the end of a trial, where they overlap with the input stimulus (Figure 1**a**-**b**). The stimulus input was also taken from an independent set of about *M* = 100 Poisson cells. Stimulus and oscillation populations were independently simulated. The stimulus lasted 0.5 seconds, and had an average firing rate of 6 Hz, and was fixed for all experiments. During the stimulus window we measured the population’s firing rate and synchrony. It is these numbers we report throughout.

**Figure 1:**
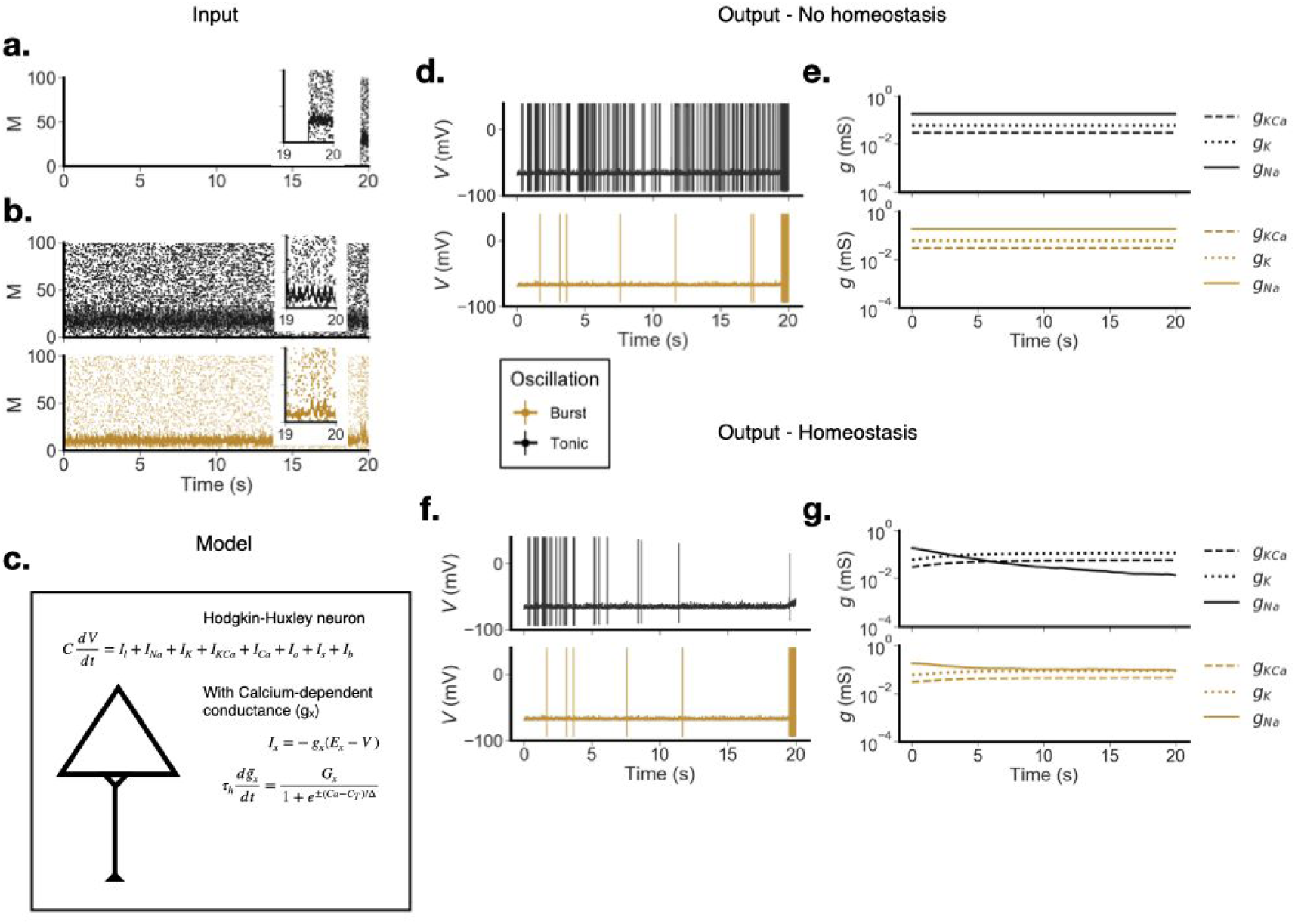
The model, its inputs, and some example results. We show examples with and without homeostasis, of both bursts (goldenrod) and tonic oscillations (black). **a**-**b**. Inputs into the model. **a.** The stimulus, which is used to estimate the model population’s synchrony and excitability. **b.** Two example input oscillations, a tonic example (black) and a burst example (goldenrod). Note: the insets in for the panels above depict input spikes over the last 1 second of run time. Black lines in these inset panels represent average firing rates calculated using 1 ms bins. **c.** Illustration of a single model neuron, its major currents, and the major equations for intrinsic homeostasis.. **d.**-**e.** Example of model output with homeostasis dynamics turned off. **d.** Is the membrane voltage, plotted in time. **e.** Depicts the Na, K, and KCa conductances, which are in this case fixed. **f.**-**g.** Example of model output, with the homeostasis dynamics turned on. **f.** Is the membrane voltage, plotted in time. **g.** Depicts changes to Na, K, and KCa conductances with time.

Each experiment began with an instantiation of a randomly generated population. A single example neuron is depicted in Figure 1**c**. This population is subjected to a range of modulatory and control conditions, including oscillatory strength, duration, and synapse type (AMPA or GABA). Each experiment lasted 20 seconds. In Figure 1**d**-**g** we depict key aspects of model output during an experiment.

After homeostatic equilibrium is reached, we measure two features: the synchrony between action potentials (measured by the Kappa correlation) and changes in the excitability of the system (measured as a change in population firing rate). Both of these measures are defined in the *Methods*. To ensure a consistent comparison between experiments, measurements were made over the same 0.5 second period in all simulations. Source code for all simulations is available at https://github.com/voytekresearch/resistingrhythm The homeostatic response in our simple model decreases the sodium conductance and increases potassium conductances (Figure 1**g**, *bottom panel*). Combined, these changes make it harder for the model to initiate action potentials and extends the recovery period after a spike. These two changes combine to explain the loss of excitability in some conditions of our model.

Note that in real systems intrinsic homeostasis is thought to happen over minutes or days. However, simulation times that are hours or days long are not computationally feasible. So we follow the field and study a model where Ca^2+^ homeostasis dynamics happen with a 4-second half-life, denoted by τ _*h*_. This might seem like a huge difference. But all that matters mathematically is that homeostasis dynamics happen much slower than all the other synaptic/membrane dynamics. A timescale of τ _*h*_ > 4 seconds is a reasonable choice to ensure this. The other major dynamics in the model are synapses with < 30 ms half-lives, leaving us with a 133-fold difference between the time scale of synaptic and homeostatic dynamics (Golowasch et al. 1999; Marder, O’Leary, and Shruti 2014; Marder, Goeritz, and Otopalik 2015; Gutierrez, O’Leary, and Marder 2013; Marder, O’Leary, and Shruti 2014; O’Leary et al. 2014; LeMasson, Marder, and Abbott 1993; Abbott and LeMasson 1993).

### Excitatory modulation

Homeostasis completely inverts the effect of tonic AMPAergic oscillatory modulation. These excitatory oscillations cause decreases in population firing rate and population synchrony. To see how, let’s consider how the population responds to a tonic excitatory modulation *without* homeostasis. Without homeostasis, increasing the strength of the excitatory oscillation increases excitability, and so increases the population firing rate (Figure 2**b**, black line).

**Figure 2:**
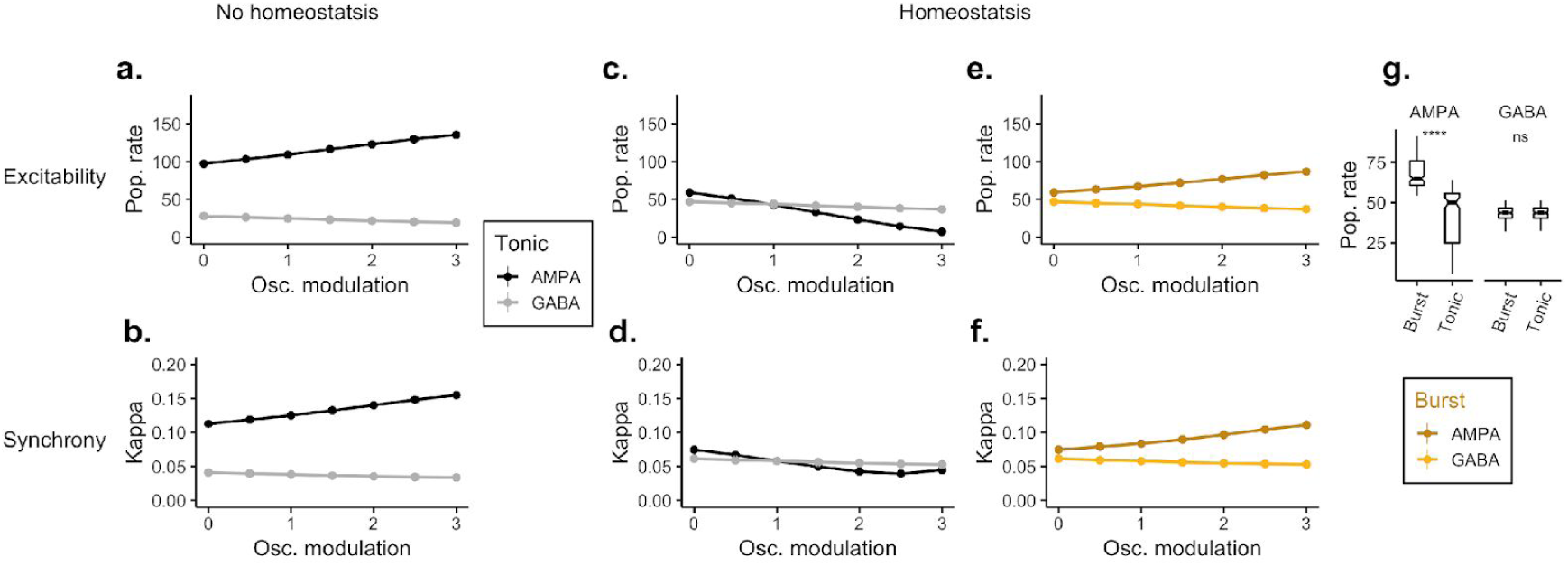
The effect of oscillatory modulation on synchrony–which is measured using the Kappa correlation (Eq. 11)–and excitability, measured by the population firing rate. Tonic oscillations are shown in grey and black. Bursts are shown in light and dark yellow. **a.**-**b.** Increases in tonic modulation strength, without homeostasis. This is our reference condition. Top panel (a.) is the observed population firing rate averaged over the 0.5 second stimulus. Bottom (b.) is synchrony over the same period. **c.**-**d.** Same experiment as a-b but with Calcium-mediated homeostasis, showing how homeostasis with tonic AMPA oscillations reduces population firing and synchrony. **e.**-**f.** Burst modulation, presented during the stimulus period (4 cycles of oscillation, onset time: 19.5 s). **g.** Change in excitability between bursts and tonic rhythms for all oscillation firing rates. Asterisks denote a significant difference using the Wilcoxon rank sum test (*W* = 1886.5, *p* < 2.2*e*−16). The frequency of the oscillatory rhythm was fixed at *f* = 8 in all models.

With homeostasis in play, this pattern inverts. As excitatory oscillatory modulation strength increases, synchrony and excitability *decrease* (Figure 2**c-d**). Homeostatic mechanisms in the model cause what *should be* excitatory synchronizing modulation to become suppressive and desynchronizing. The stronger the oscillation, the more suppressive the result. By the time the oscillation is about half the strength of the stimulus (which we fix at a firing rate of 6 Hz) the response to the stimulus is completely suppressed and the population firing rate approaches 0 (Figure 2**c**).

We compared the effect of these tonic oscillations to short 3-cycle bursts of modulation, presented only during the stimulus. Here, the oscillation period is far too short to provoke much, if any, homeostasis. This lets bursts sidestep our early results, and so increases in oscillatory strength continue to increase population firing rate and synchrony (Figure 2**e** and **f**).

Of course theta oscillations in the hippocampus are GABAergic (Colgin 2016a), not AMPAergic as we model here. However by exploring artificial AMPAergic oscillation we can demonstrate the central principle we want to highlight. It seems that tonic oscillations–when joined with homeostasis–can profoundly alter the coding properties of what one would typically expect from oscillatory modulation, and the degree of this effect depends entirely on the degree to which the oscillation changes the cell’s tonic Ca ^2+^. And that because of this, bursts of oscillation are *qualitatively distinct* from their tonic counterparts.

The specific effect we observed (suppression of excitability) is seen because the homeostatic equations that *we chose* respond to changes in Ca^2+^ by decreasing the conductance of the Na, and increasing the conductance of K and KCa channels (Eq. 8). This makes firing an action potential less likely, for any given input. These channel dynamics are visualized in Figure 1**g** and a corresponding FI curve is shown in Figure 4.

**Figure 3:**
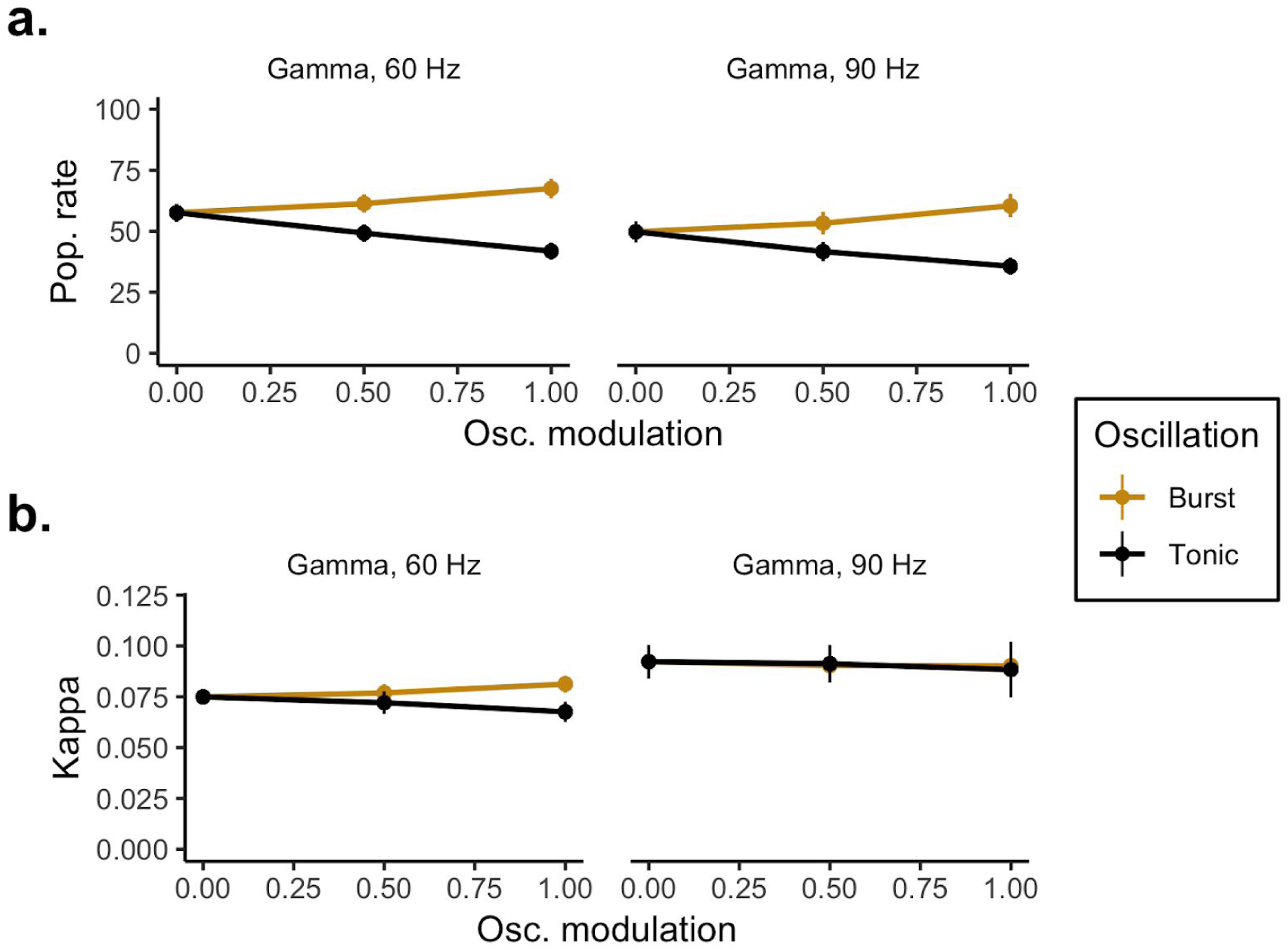
The effect of fast AMPAergic gamma oscillations for both tonic oscillation and 3-cycle bursts. Top panel (**a**.) is the observed population firing rate averaged over a 0.05 second or 0.033 second stimulus. Bottom (**b**.) is synchrony over the same period. Columns in both a. and b. represent with 60 or 90 Hz gamma. The stimulus window was matched to the length of time 3 cycles of oscillation would take. To generate this curve the model was equilibrated for 20 seconds, under a 1.5 Hz firing rate and 8 Hz frequency rhythm, after which the current impulse that we described above was applied. During FI-curve estimation there was no oscillation present.

**Figure 4:**
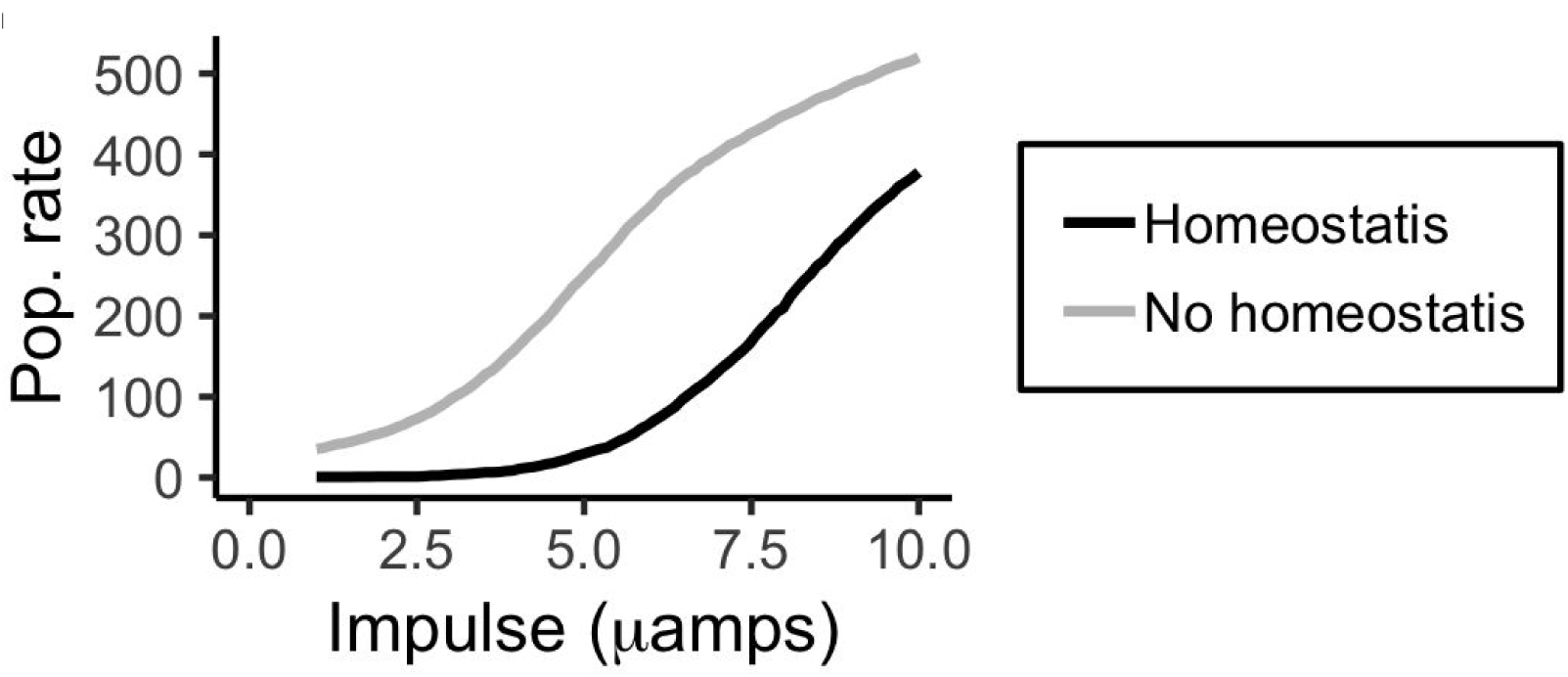
Excitability and homeostasis. An average FI-curve for 100 hippocampal neurons, with (black) and without (grey) homeostasis. In both models, oscillation was tonic and AMPAergic. The square-wave pulse lasted 0.1 seconds, and ranged from 0.1 to 1 μ amps, sampled every 0.01 μ amps.

The general effect we would expect for some other selection of channels, chosen by some other theorists, or by evolution itself, depends strongly on those selected channels. O’Leary (O’Leary et al. 2014) has made it clear how complex the relationship is between homeostasis, channel conductance, and firing patterns. It’s so complex that we believe general statements are probably impossible, at least as they might relate to this paper. While our simple model has been shown by others to recapitulate hippocampal pyramidal cell’s activity in laboratory settings (LeMasson, Marder, and Abbott 1993), it is a *very simple* model, and validating our work does require new experimental testing. This could be done, for example, by using prolonged excitatory gamma oscillations–arising from direct stimulation of either the medial entorhinal cortex or CA3–to drive a possible homeostatic response in CA1 (Csicsvari et al. 2003; Colgin et al. 2009).

Our model shows that there is sometimes an inversion in population synchrony, as seen in Figure 2**d** and 5**b**. We believe this is a low *N* effect: as population spiking is suppressed, the total number of spikes declines to the point where the bins used to calculate the Kappa correlation are often empty. This leads to a somewhat misleading “increase” in synchrony.

### Inhibitory modulation

GABAergic oscillatory modulation does not lead to a homeostatic response in our model. This is true for both tonic oscillations and bursts. No matter the homeostatic state, as GABAergic strength increases population firing declines (grey lines in Figure 2**a-d** and light yellow lines in panel **e-f**).

In the work so far, Figure 2, we compared AMPA and GABAergic models of the theta rhythm (8 Hz). This was done to illuminate their different effects, but was not a biologically realistic model. Hippocampal theta is solely a GABAergic rhythm, at least when driven by medial septum (Colgin 2016). AMPAergic oscillations are, however, seen in the high gamma range (60-90 Hz) (Csicsvari et al. 2003; Colgin 2016a). In Figure 3 we confirm AMPAergic oscillations in the gamma range follow the same pattern as theta. Both 60 and 90 Hz gamma rhythms can suppress excitability and desynchronize firing.

### Homeostasis and excitability

In our model Ca^2+^ influx is driven by increases in the membrane voltage but the Ca^2+^ concentration is restored by passive and linear recovery (Eq. 9), and active homeostasis (Eq. 8). Homeostasis works by negative feedback. Increases in Ca^2+^ concentration decrease the conductance of both the Na and K channels in our model (Eq. 7). This decrease in conductance in turn lowers the membrane voltage down closer to its resting value. As the voltage falls so does the Ca^2+^ concentration. The excitability changes we see with homeostasis must be due to the decreases in Na conductance (which initiates spikes) and increases in K conductance (which recovers the voltage following a spike).

One of the simplest, most classic ways to measure the excitability of a cell–simulated or real–is with an FI-curve. In an FI-curve a small square wave of current is injected into the cell, and the resulting increase in firing rate (if any) is averaged over a short window. In Figure 4 we show the average FI-curve for 100 simulated hippocampal neurons. Under homeostasis the FI-curve is lower, which means there has been a substantial suppression in the membrane excitability.

### Controls and perturbations

To ensure our parameter choices in the model were generalizable, we perturbed several key values by 10-20 percent.

When the model is run with only stimulus-driven homeostasis, the Ca^2+^ concentration equilibrates to about 0.003 mM. We used this as a standard target value for all modulation experiments, until now. When we vary this value in 0.0002 mM increments, population rate and synchrony either increases or decreases depending on whether the Ca^2+^ increases or decreases, shown Figure 5**a-b**. However despite different initial Ca^2+^ concentrations, each model still shows an identical set of trends as the strength of the oscillation increases (Figure 5). That is, increasing or decreasing the target concentration shifts the overall excitability of the population, in an approximately linear way. This means that while the initial choice of 0.003 mM was arbitrary, the qualitative pattern of results we report is not dependent on this choice.

**Figure 5:**
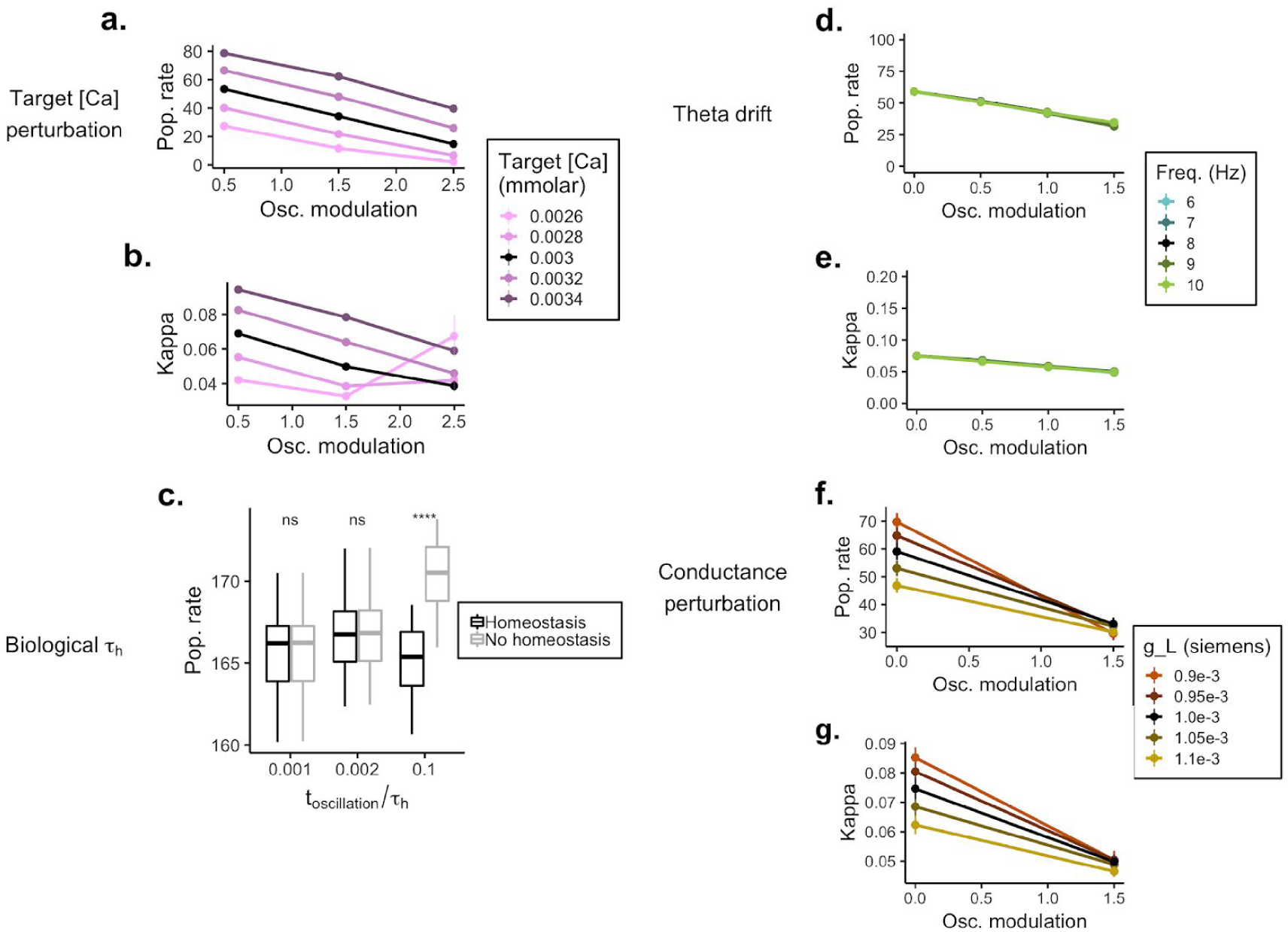
Control experiments. **a.** Population firing rate for different levels of target [Ca] (colors). All values are referenced to a no-modulation control. **b.** Change in population synchrony for different levels of target [Ca] (colors). All values are referenced to a no-modulation control. **c.** Change in population rate as the oscillation duration approaches a more realistic τ _*h*_, the half-life of the homeostasis dynamics (Eq. 8). In this control experiment we used a more biologically realistic τ _*h*_ of 600 s (or 10 minutes). All other simulations in the report use a τ _*h*_ of 4 seconds, which is well below most reports of this value in real systems. However in choosing such a small value we follow the majority of the homeostasis modeling literature (for more on this see the Discussion). **d**.-**e**. The effect of frequency drift (see legend) on homeostasis with tonic AMPAergic modulation. **f**.-**g**. The effect of altering the leak conductance (*g*_*L*_) on homeostasis with tonic AMPAergic modulation.

We chose a 3-cycle length for bursts somewhat arbitrarily. Real oscillations range in duration from single cycles to seconds of even minutes (Lundqvist et al. 2016; van Ede et al. 2018). Our burst/tonic distinction is therefore a false (but useful) dichotomy. To find a better estimate of how long an oscillation would need to last, on average, to generate a homeostatic response we ran some long simulations using a more realistic homeostatic time constant (τ _*h*_ = 600 seconds; up from 4 seconds). It seems an oscillation would need to be present *at least* 10% of the time to generate a response (Figure 5**c**). Jacobs (Jacobs 2014) estimated the prevalence of hippocampal theta to be about 40% of the time in rats, and 10% in humans. So even for humans, who have slower and less stable theta rhythm(s) (Goyal et al. 2018), we’d expect an homeostatic response from AMPAergic theta, if it existed.

Real theta rhythms drift in frequency (Buzsáki and Schomburg 2015; Jacobs 2014). To test if small differences in frequency have a notable effect, we simulated small drifts in frequency of theta ranging from 6-10 Hz. Frequency drift had no visible effect on the homeostatic suppression of excitability (Figure 5**d**-**e**).

The leak conductance (*g*_*L*_, Eq. 7) plays a key role in setting the resting potential of the cell. The resting potential in turn helps determine the overall membrane excitability, and sets the resting Ca^2+^ concentration. This means our Ca^2+^ set point (of 0.003 mM) depends on *g*_*L*_. To test if a small mismatch between *g*_*L*_ and the Ca^2+^ set-point altered our results, we ran several experiments perturbing the leak conductance while keeping Ca^2+^ fixed, but in the end mismatch in leak conductance had only a small effect on homeostasis (Figure 5**f**-**g**).

## Discussion

Our scientific understanding of homeostasis has been shaped as much by theoretical work as empirical (Marder, O’Leary, and Shruti 2014). In an attempt to understand the interaction between oscillatory modulations and homeostasis, we began by studying one of the simplest models used in early studies of homeostasis–a population of point neurons in the hippocampus (LeMasson, Marder, and Abbott 1993).

Our simple models offer answers to three basic questions. One, do tonic oscillations engage homeostatic mechanisms? Two, does homeostasis in turn change the oscillation’s function? Three, do short bursts of oscillation have distinct effects from tonic oscillations? Put another way: can homeostasis explain why some oscillations, such as hippocampal theta, tend to appear as tonic rhythms while other oscillations (like those in cortex) tend to appear as bursts? (Peterson and Voytek 2017; Cole and Voytek 2018; van Ede et al. 2018)

Oscillations are generally conceived to play a functional role in modulating spiking (Fries 2015; van Ede et al. 2018) by grouping action potentials and improving signal to noise (Chen et al. 2013; Zhou et al. 2015; Voytek et al. 2015; Peterson and Voytek 2017). Absent the effects of homeostasis, our model clearly demonstrates this: with increasing modulation strength, excitatory oscillations show increased firing rate and synchrony. Given this effect, and the importance of phase coding in the hippocampus (Lisman and Jensen 2013), it is reasonable to wonder why hippocampal oscillations are GABAergic. Our model suggests that while this is true for brief, bursty excitatory oscillations, stronger, tonic excitatory oscillations engage homeostatic mechanisms that alter Na and K conductances, driving down the membrane voltage. This, in turn, reduces both the firing rate and spike synchrony in response to inputs.

Homeostasis is generally viewed as a stabilizing force. This is the goal of homeostasis in our model as well. However oscillatory modulation is a unique form of modulation; it shares neurotransmitters, and even synapses, with non-modulatory stimuli. As a result, homeostatic corrections for modulation must affect both the modulator and the driver (Sherman and Guillery 1998). Our work suggests this trade-off can be quite strong. To avoid this trade-off, sometimes anyway, we speculate that the long-time scale of evolution may sometimes favor different types of oscillation for different functional roles.

Bursting, rather than sustained, oscillations tend to be common in the cortex. One striking example is in motor cortical regions, where beta (12-30 Hz) bursts are prevalent and likely functional. Specifically, beta bursts are very short–sometimes lasting only one or two cycles (Sherman et al. 2016; Cole and Voytek 2019; Jones 2016)–and relate to self-timed movements (Feingold et al. 2015). Patients with Parkinson’s disease show increased motor rigidity and bradykinesia, symptoms associated with prolongation of beta bursts (Tinkhauser et al. 2017). Levodopa treatment was shown to decrease burst probability and duration, and that decrease in burst duration correlated with motor improvement (Tinkhauser et al. 2017). Our work here suggests some of the pathophysiology in Parkinson’s may be due to intrinsic homeostasis.

An important counterexample to healthy beta bursts is cortical alpha (8-10 Hz) in visual cortex. Alpha is tonic and high powered when humans rest with their eyes closed. Even though both cortical and subcortical alpha generators are synaptically excitatory, this rhythm has however been associated with suppression of excitability (Jensen 2002; Bonnefond and Jensen 2012; Peterson and Voytek 2017). While several competing explanations have been offered for this (Bonnefond and Jensen 2012; Lange, Oostenveld, and Fries 2013; Peterson and Voytek 2017), our work raises another possibility that is complementary to the others. Specifically, though we modeled hippocampal cells, the same Ca^2+^ homeostatic mechanisms exist in visual cortex (and the neocortex as a whole). This means that strong and tonic alpha oscillations, combined with homeostasis, should directly suppress population firing in visual cortex. Such an effect, were it to occur, would last well past the moment of oscillation offset an effect of alpha that has been reported in the literature (Jensen 2002; Bonnefond and Jensen 2012). In these studies the physiological mechanism was unclear. Our work suggests that it might be intrinsic homeostasis.

## Limitations

The nature of our model–the fact that we use point neurons with only 6 currents–or the fact that our model is strictly feed-forward–without lateral or recurrent connections–means we don’t know with confidence to what degree our model’s effects will appear in more complex models, or in real neural systems. O’Leary (O’Leary et al. 2014) and others have shown how exceptionally complex the homeostatic response is. It is therefore likely some neurons will have a homeostatic response that is nothing like that in our model. Others may be naturally immune to oscillatory effects. There may be many such examples.

We do know, however, that oscillations are a ubiquitous feature of cortical and subcortical activity, as is Ca^2+^-mediated intrinsic homeostasis. This means the ingredients for oscillation and homeostasis to interact are omnipresent in both subcortical and cortical areas. This response might be very different from what we found.

The basic mechanisms of Ca^2+^-mediated intrinsic homeostasis are conserved across phyla (Tran et al. 2017). Meaning the qualitative properties of our model may also be present across phyla. That said, the mechanisms for homeostatic interactions, along with their quantitative properties, depend on a number of factors specific to each cell and circuit. These include the duty cycle, power, and frequency of an oscillation, as well as on synaptic strengths and their location in the dendritic tree (an idea we return to below). It also depends on the other inputs into the cell, both from fast synaptic transmission and other (slower) modulators, as well as connection type; simulation studies suggest that recurrent connections can strongly interact with homeostatic regulation (Harnack et al. 2015). The temporal and spatial scales of these factors will strongly influence Ca^2+^ dynamics, which is in turn central to governing what, if any, homeostatic effects oscillations may generate.

We did not study synaptic homeostasis. Synaptic homeostasis can also play a key role in combating oscillatory perturbations (Cannon and Miller 2016). There is some evidence that intrinsic homeostasis appears linked to excitatory synaptic homeostasis (Joseph and Turrigiano 2017). Still, the exact nature of the homeostatic response depends on how, and to what degree, oscillatory and other sensory or internally driven inputs share synapses. This, in turn, requires considering complex dendritic arbors and their effect on homeostasis (LeMasson, Marder, and Abbott 1993), neuromodulation (Jadi et al. 2012; Jadi and Sejnowski 2014), and computation (Mainen and Sejnowski 1996; Polsky, Mel, and Schiller 2004; Mel and Schiller 2004). Considering these interacting effects together is the next step needed to develop a clearer biological understanding of modulatory oscillations. For inhibitory oscillations the case is even more complex: Ca^2+^ in these cells does not appear to regulate intrinsic homeostasis, but synaptic homeostasis is under a separate mechanism of control (Joseph and Turrigiano 2017).

### Previous work

Homeostasis has been extensively studied in the rhythmic pacemaker present in the crab and lobster stomatogastric ganglion, where homeostasis stabilizes these oscillations (Golowasch et al. 1999), and interacts with neuromodulation in a *highly* state dependent way (Marder, O’Leary, and Shruti 2014; Marder, Goeritz, and Otopalik 2015; Marder, O’Leary, and Shruti 2014). Like our model though, the oscillation present in the stomatogastric ganglion is tonic. This means it could in principle drive a secondary homeostatic response. However, synapses in the stomatogastric ganglion circuit are predominantly inhibitory (Marder, Goeritz, and Otopalik 2015). If our model predictions held up for this circuit we would expect there to be no secondary response.

The interaction between homeostasis and oscillations has previously been considered in a simple model of cortex. Cannon and Miller (Cannon and Miller 2017) explored how synaptic homeostasis can effectively minimize the effect of modulatory perturbations, thus maximizing mutual information between an incoming oscillatory signal and a single cell’s firing pattern. But the oscillatory input in this work was treated as a signal, and not a modulator itself (Cannon and Miller 2017). We believe this is an important distinction, and our analysis is an inverse complement to (Cannon and Miller 2017). We studied how to minimize the perturbation caused by an oscillator, rather than how to maximize its transmission.

## Conclusion

Here, using a relatively simple model of hippocampal neurons, we observe a surprising–even paradoxical–result: that homeostatic effects can invert a normally synchronizing excitatory oscillatory neuromodulator and cause it to become inhibitory and desynchronizing.

If our simple model is right, we conjecture that intrinsic homeostasis can explain why tonic theta rhythms in the hippocampus are synaptically inhibitory. To make this clearer, consider the alternative: if the theta rhythm was strong, tonic, and synaptically excitatory, our model suggests this could lead to an equally strong–but opposing–homeostatic response. Such a response can mean that sodium conductance decreases, which in turn decreases the likelihood a hippocampal neuron would respond to any given stimulation.

In effect, the homeostatic response to strong excitatory oscillations consumes a substantial portion of each cell’s dynamic range. On the other hand, inhibitory oscillations–which do not generate an intrinsic homeostatic response–leave the dynamic range of neurons intact. This may also explain why neocortical oscillations tend to be short and bursty, and why some neurological disorders, such as Parkinson’s disease, are associated with prolonged excitatory rhythms.

## Methods

### Mathematical model

We model an unconnected population of hippocampal neurons, subjected to oscillatory modulation. This is instantiated as *N* = 1000 input cells connected to *M* = 100 Hodgkin-Huxley neurons. The *M* cells in the population were tuned to mimic regular firing (Borgers, Epstein, and Kopell 2005; Borgers, Epstein, and Kopell 2008). The firing pattern of each input cell (both stimulus and modulation) is from a Poisson process, with a time-varying rate. *N*_*o*_ cells oscillate. *N*_*s*_ serve as input. For simplicity, we let 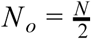 so *N*_*o*_ = *N*_*s*_. All input cells have a *p* = 0.1 connection probability to the hippocampal population. The hippocampal population has no lateral or recurrent connections. The synaptic weights for all *N* → *M* connections *w* were independently sampled from a uniform distribution, *w* ∼ *U* (5, 50) μ S. The firing rate of the oscillating population was governed by a biased sinusoidal pacemaker, with amplitude 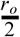 and frequency *f*, with the exact form 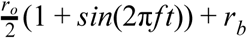. As a result, *r*_*o*_ defines the peak firing of the oscillation but *r*_*o*_ + *r*_*b*_ is the total drive experienced by each cell. For simplicity though we remove the constant bias term *r*_*b*_ in our plots (Figures 2,3,5); it is present in all our simulations, tonic oscillations and bursts. The stimulus population was simply modeled by a fixed rate of 6 Hz (*r*_*s*_).

Hodgkin-Huxley dynamics were governed by 4 active ionic currents (*I*_*Na*_, *I*_*K*_, *I*_*KCa*_ *I*_*Ca*_) and the passive leak current (*I*_*l*_ = *g*_*l*_(*E*_*l*_−*V*)). Besides *I* _*Ca*_ (which is discussed below), active currents are governed by the standard Hodgkin and Huxley form (Hodgkin and Huxley 1952). Where *m* and *h* respectively track the opening and closing channel kinetics, *p* and *q* are channel dependent parameters, and *V*_*i*_ is the channel appropriate Nernst reversal potential. See *Table 1* for the complete set of parameters.

**Table 1:** Model parameters.

| Symbol | Range (unit) | Description |
| --- | --- | --- |
| $f$ | 8 (Hz) | Oscillation frequency |
| $r_o$ | 0 - 6 (Hz) | Oscillation firing rate |
| $r_s$ | 6 (Hz) | Stimulus firing rate |
| $r_b$ | 2 (Hz) | The “background” firing rate |
| $C$ | 1 ($\mu$ F) | Membrane capacitance |
| $\tau_h$ | >4 (sec) | Homeostasis time constant |
| $w$ | 5 - 50 ($\mu$ S) | Synaptic weight |
| $\tau_e$ | 5 (msec) | Excitatory synaptic time constant |
| $\tau_i$ | 10 (msec) | Inhibitory synaptic time constant |
| $V_e$ | 0 (mV) | Excitatory synaptic reversal potential |
| $V_i$ | -80 (mV) | Inhibitory synaptic reversal potential |
| $p$ | 0.1 | Connection probability (stimulus, modulation) |
| $C_T$ | 0.0028-0.0032 (mM) | Target $\text{Ca}^{2+}$ concentration |
| $G_{Na}$ | 360 ($\mu\text{S}$) | Initial Na conductance |
| $G_K$ | 120 ($\mu\text{S}$) | Initial K conductance |
| $G_{KCa}$ | 60 ($\mu\text{S}$) | KCa conductance |
| $g_{Na}$ | 180 ($\mu\text{S}$) | Max. Na conductance |
| $g_K$ | 60 ($\mu\text{S}$) | Max. K conductance |
| $g_{KCa}$ | 30 ($\mu\text{S}$) | Max. K conductance |
| $g_{Ca}$ | 0.03 ($\mu\text{S}$) | Max. $\text{Ca}^{2+}$ conductance |
| $g_l$ | 1 ($\mu\text{S}$) | Leak conductance |
| $V_l$ | -70 (mV) | Leak reversal potential |
| $V_K$ | -100 (mV) | K reversal potential |
| $V_{Na}$ | 50 (mV) | Na reversal potential |
| $V_{Ca}$ | 150 (mV) | $\text{Ca}^{2+}$ reversal potential |
| $V_1$ | -50 (mV) | Morris-Lecar constant |
| $V_2$ | 10 (mV) | Morris-Lecar constant |
| $\Delta$ | 0.6 ($\mu\text{M}$) | $\text{Ca}^{2+}$ influx rate |
| $k$ | 1/200 (1 / msec) | $\text{Ca}^{2+}$ clearance rate |
| $\gamma$ | -0.00047 (mM / mA / msec) | $\text{Ca}^{2+}$ current conversion constant |
| $\delta t$ | 0.01 (msec) | Integration time step |

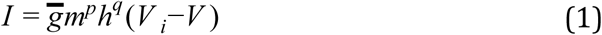

The Ca^2+^ current *I*_*Ca*_ was governed by a form taken from the Morris-Lecar model (Morris and Lecar 1981; LeMasson, Marder, and Abbott 1993; Siegel, Marder, and Abbott 1994).

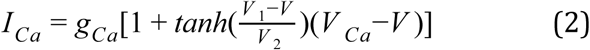

Overall membrane dynamics were governed by these internal ion conductances, and two synaptic input terms, *I*_*s*_ and *I*_*o*_. These are, respectively, the stimulus and oscillatory inputs. All synaptic input, both stimulus and oscillation were governed by single exponential kinetics, which we denote generically using an *x* subscript below.

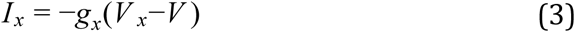

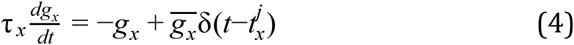

As a concrete example of using the abstract form in Eq. 3-4, let’s define the stimulus. Spikes arriving from some jth neuron are denoted by *t*_*x*_^*j*^. The stimulus was always excitatory, so to define *I*_*s*_ we substitute in AMPAergic voltage, conductance, and time constant values taken from Table 1 in for *x* (above), giving Eqs. 5-6.

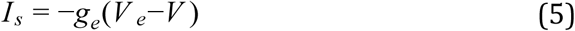

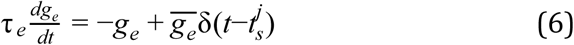

To define *I*_*o*_ we make a similar set of substitutions, choosing either AMPAergic or GABAergic values, depending on which kind of oscillation we wish to simulate. Having defined both internal dynamics and synaptic dynamics we arrive at the final model, stated in terms of its biophysical currents (Eq. 7).

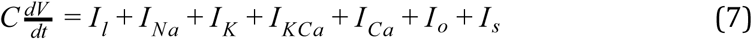

Eq. 7 would mark the end of a traditional Hodgkin-Huxley definition. However modelling intrinsic homeostasis means altering both inward and outward conductances in response to changes in Ca^2+^ concentration. Following the previous work of (LeMasson, Marder, and Abbott 1993) and (Siegel, Marder, and Abbott 1994), we modeled this by having maximal conductances 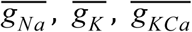, and 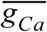 non-linearly vary with the Ca^2+^ concentration, *Ca* (Eq.8). In this scheme and during homeostatic equilibration, conductances drift until the target Ca^2+^ concentration is met, denoted as *C*_*T*_. In a control experiment a range of *C*_*T*_ values were explored (Figure 5), though simulations default to 0.003 mM; the value the system reaches with stimulation (*r*_*s*_ = 6 Hz) without modulation (*r*_*o*_ = 0). The ± symbol in equation 8 denotes the direction of ion flow and is (+) for inward going currents and (−) for outward going.

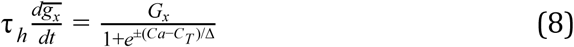

Ca^2+^ dynamics were assumed to follow first order kinetics, driven by the Ca^2+^ influx current and clearance rate constant *k*. Values for both γ and *k* were taken from (Liu et al. 1998).

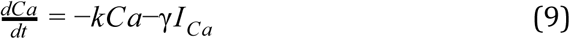

### Estimating excitability and synchrony

We measure excitability by comparing average firing rate of all *M* neurons in an experiment, with (*r*_*m*_) and without modulation 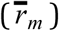.

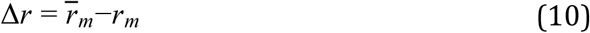

We measure synchrony using κ, a binned measure of spiking covariance (Wang and Buzsáki 1996). Where *X* (*l*) = 0 *or* 1 and *Y* (*l*) = 0 *or* 1 for *l* = {1, 2, …, *K*} and with *T* /*K* = τ.

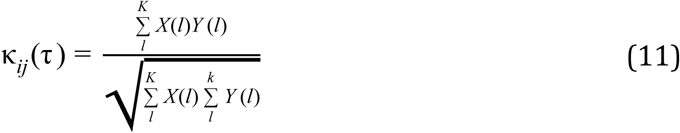

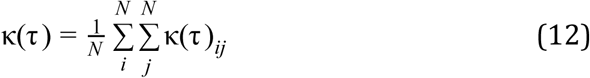

